# Protein Hunter: exploiting structure hallucination within diffusion for protein design

**DOI:** 10.1101/2025.10.10.681530

**Authors:** Yehlin Cho, Griffin Rangel, Gaurav Bhardwaj, Sergey Ovchinnikov

## Abstract

Interactions between proteins and other biomolecules underlie nearly all biological processes, yet designing such interactions de novo remains challenging. Capturing their specific interactions and co-optimizing sequence and structure are difficult and often require extensive computation. We present Protein Hunter, a fast, fine-tuning-free framework for de novo protein design. Starting from an all-X sequence, we find diffusion-based structure prediction models hallucinate reasonable looking structures that can be further improved through iterative sequence re-design and structure re-prediction. This lightweight strategy achieves high AlphaFold3 in silico success rates across both unconditional and conditional generation tasks, including binders to proteins, cyclic peptides, small molecules, DNA, and RNA. Protein Hunter also supports multi-motif scaffolding and partial redesign, providing a general and efficient platform for de novo protein design across diverse molecular targets.

## 2 Introduction

Deep learning and computational methods have revolutionized protein design, enabling de novo creation of binders, catalysts, and functional assemblies for applications ranging from antibody therapeutics to industrial enzymes [Kortemme, 2024, Pan and Kortemme, 2021].

Current protein design methods broadly fall into two categories: generative and optimization-based. Generative diffusion models like RFdiffusion Watson et al. [2023], Ahern et al. [2025], Butcher et al. [2025], Vázquez Torres et al. [2024] can create novel protein folds, assemblies, and binders for diverse targets including proteins, small molecules, and nucleic acids. However, its sequential structure-then-sequence workflow cannot guarantee that final designs satisfy both structural and sequence constraints simultaneously, as the generated backbones may be suboptimal for designing sequences that fold into the intended structure [Frank et al., 2024, Cho et al.]. This requires generation of large number of designs followed by filtering. Optimization-based methods such as BindCraft Pacesa et al. [2025] and BoltzDesign [Cho et al., 2025] address this by iteratively co-refining structure and sequence, yet their reliance on gradient decent leads to slow convergence and susceptibility to local minima. Both approaches face prohibitive computational costs for large or multimeric designs due to large GPU memory requirements.

Recent advances in all-atom structure prediction suggest an alternative strategy. AlphaFold3 [Abramson et al., 2024, Jumper et al., 2021] and similar models like Boltz-1/2 Passaro et al. [2025], Wohlwend et al. [2025] and Chai-1 team et al. [2024] use diffusion-based approaches for all-atom structure prediction. Similar to how diffusion models hallucinate beyond training distributions in image generation [Aithal et al., 2024, Cohen et al., 2024], we found these models can hallucinate high-quality folded structures even from sequences that would not naturally adopt those conformations by using learned structural priors to denoise toward well-folded backbones. Rather than treating this as a defect, we repurpose it for protein design.

Here, we present Protein Hunter, a fast, fine-tuning-free framework that exploits this capability through iterative structure-sequence cycling. Starting from a sequence of unknown tokens (all-X), an AlphaFold3-style models hallucinates an initial fold, which is then refined through cycles of sequence design with ProteinMPNN and structure prediction. The approach is efficient—requiring 10 seconds for a 100-residue protein and 130 seconds for a 900-residue design. Protein Hunter supports unconditional design, binder design for proteins, cyclic peptides, small molecules, and nucleic acids, as well as motif scaffolding and partial redesign.

## 3 Method

Our method leverages the ability of diffusion-based structure prediction models to hallucinate protein-like folds from out-of-distribution inputs. To quantify this, we tested single-amino acid repeats and compared their distograms with natural protein distance distributions (see Appendix).

### 3.1 Unconditional and Conditional Protein Design

To evaluate the ability of diffusion-based structure prediction models to hallucinate realistic protein backbones, we performed unconditional protein generation across sequence lengths of approximately 100–900 residues using AlphaFold3 (AF3)-style models, including Chai-1, Boltz-2, and AF3. Each position was initialized with the unknown X token, and 20 sequences per length were generated. For each resulting structure, a single sequence was designed using ProteinMPNN/LigandMPNN, refolded with AF3, and assessed by backbone RMSD (pre-vs. post-refolding) and pLDDT to measure structural consistency and confidence. For conditional protein generation, relevant to binder-design tasks such as the BindCraft Pacesa et al. [2025] benchmarks, the target sequence and multiple sequence alignment (MSA) were provided as inputs. Binder residues were initialized entirely with X tokens, and the model iteratively generated and refined the sequence to obtain high-confidence designs.

### 3.2 Iterative Structure-Sequence Cycling with Protein Hunter

Initial structures generated by diffusion hallucination are often imperfect, lacking side-chain information and potentially incomplete folds. ProteinMPNN can also fail when provided with noisy or poorly folded backbones [Cho et al.]. As demonstrated by AF2cycler and LASErMPNN[Frank et al., 2024, Fry et al., 2025], cycling between structure prediction and sequence redesign overcomes these limitations. Structure prediction refines the backbone toward conformations favored by the sequence model, while sequence redesign adapts amino acid composition to the predicted backbone, iteratively improving foldability and designability.

We implemented an iterative structure-sequence cycling framework initialized by diffusion hallucination (Figure 1D). Our method alternates zero-shot structure prediction with ProteinMPNN-based sequence redesign to progressively guide the backbone toward more designable conformations. textbfInitialization: Start with an all-X (or UNK) sequence, with or without target information, and predict its structure using an AF3-style model. **Sequence Redesign**: Redesign the sequence using ProteinMPNN (or LigandMPNN for ligand-binding tasks) and re-predict the complex structure. **Iteration** : Repeat the redesign and prediction steps for *N* iterations or until convergence.

**Figure 1:**
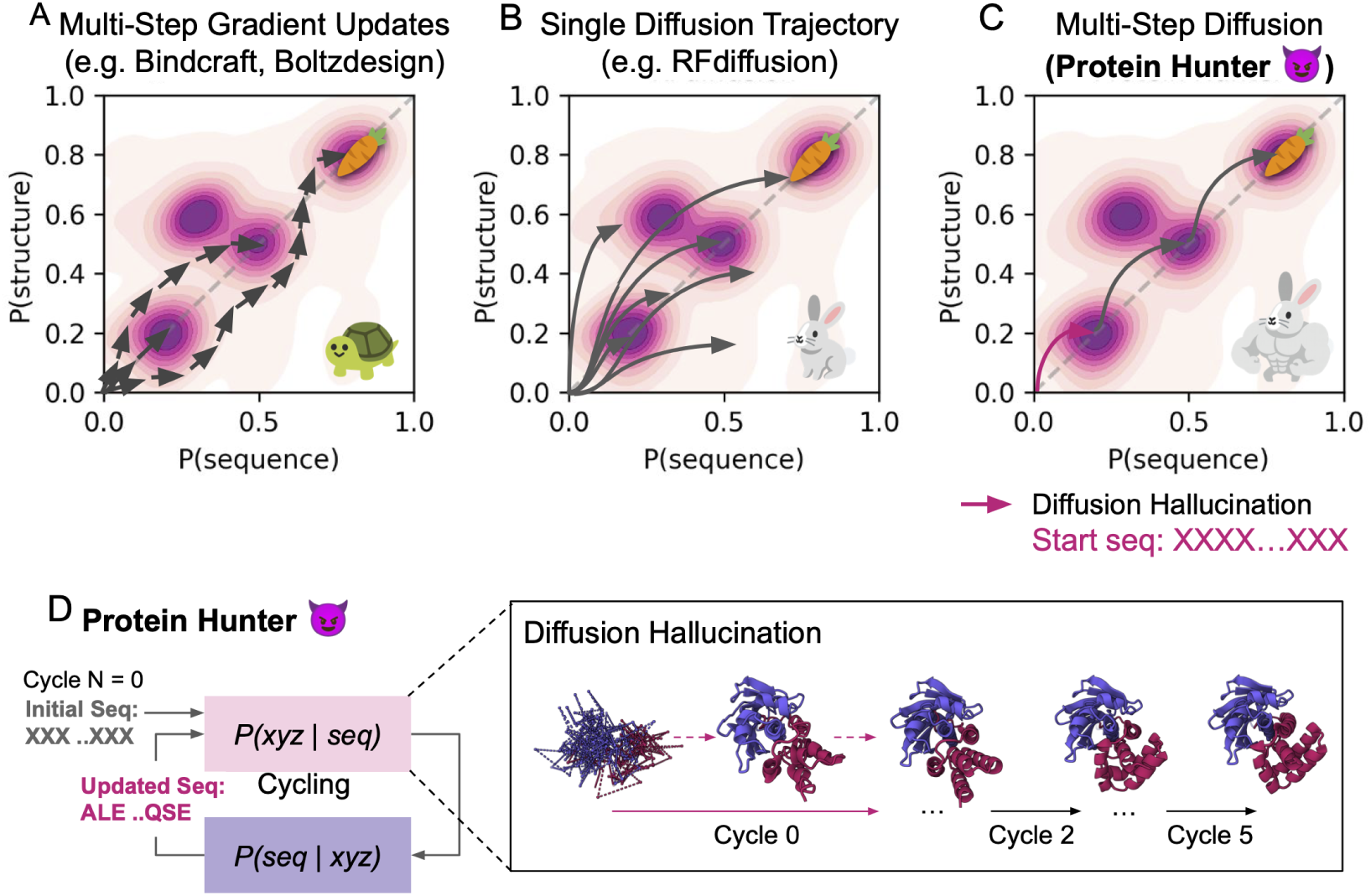
Scheme comparing sequence–structure optimization processes across different protein design models: (A) Multi-step, gradient-based methods (e.g., BindCraft, BoltzDesign). (B) Single diffusion trajectory approaches (e.g., RFdiffusion). (C) Multi-step diffusion methods (e.g., Protein Hunter) that use diffusion hallucination to jointly optimize sequence and structure. (D) Protein Hunter cycles between structure prediction and sequence design; shown is binder generation for the Bet v I allergen at cycles 0, 2, and 5.

For cycler designs, we generated one sequence per cycle and evaluated its success, rather than sampling multiple sequences, reflecting that the cycling process itself optimizes sequence space.^2^ Unlike AF2cycler, with limited hallucination ability in the structure module, we find Protein Hunter does not require a starting point and the sequence can be replaced completely at each iteration.

## 4 Results

### 4.1 Protein Hunter Generates Protein Backbones Using the X Token via Diffusion Hallucination

We first evaluated how sequences composed of a single amino acid differ in their propensity to hallucinate and generate designable protein structures. Sequences of histidine (H) or asparagine (N) showed the highest Spearman correlations (∼ 0.6) with the background residue–distance distribution, followed by the unknown X token (∼ 0.5) (Fig. 4A). To further assess their potential for binder design, we performed conditional generation using Boltz-2 and sequence redesign with SolubleMPNN, starting from single–amino-acid sequences [Passaro et al., 2025]. Among all tokens, X-initiated designs achieved the highest ipTM scores and the lowest loop fractions, indicating the most compact and stable folds (Fig. 4B,C). AF3-style models encode unknown PDB residues as X tokens and explicitly tokenize non-canonical amino acids, D-amino acids, and ligand atoms. This enables folding of sequences containing unspecified or unconventional residues while minimizing bias from amino acid–specific features.

Across diffusion-based structure prediction models, Boltz-2 achieved the highest in silico success rate (Fig. 7A), with high structural confidence and low RMSD between original and redesigned structures. We hypothesize that the strong performance of Boltz-2 arises from two key features. First, Boltz-2 encodes each non-canonical residue as a single learned token, simplifying novel chemistry and allowing the X token to act as a single residue. In contrast, AF3 and Chai-1 use atom-level tokenization, which increases sampling complexity. Second, Boltz-2 applies rigid-body alignment at every reverse diffusion step constraining the denoising path and reducing inference drift while preserving structural compactness. AF3 reduces hallucination partly through fine-tuning on a distillation predictions from AF2, which in turn encourages spaghetti-like structures for low-confident regions [Abramson et al., 2024]. In contrast, Boltz-2 and Chai-1, which were not fine-tuned on AF2 predictions to preserve the spaghetti-like structure, conveniently hallucinate ordered, well-packed structures in low-confidence regions.

Finally, structures generated by AF3-style models exhibited lower structural diversity than those from RFdiffusion (Fig. 7C), reflecting a bias toward *α*-helical topologies. This helical preference, common in generative models [Dawson et al., 2017, Lin, 2024], constrains fold diversity. To enrich *β*-sheet content, we introduced a negative helix bias into the Pairformer pair features prior to diffusion, which increased the proportion of sheet-containing samples (Fig. 12).

### 4.2 All-Atom Protein Binder Design via Structure Prediction and Sequence Cycling Enables High In Silico Success Rates

Similar to AF2cycler [Frank et al., 2024], Protein Hunter cycling structure–prediction model achieves high designability and high AF3 success rate. Initial backbones generated from the X token were often redesigned with high alanine content by ProteinMPNN, since X tokens share alanine’s backbone atoms (N, C_*α*_, C, C_*β*_). Repeated structure–sequence cycles refine overall geometry, improve secondary-structure packing, reduce alanine bias, and yield more stable folds (Fig. 9A).

During unconditional generation, single-pass designs longer than ∼300 residues often exhibited pLDDT scores below 0.7, whereas cycling maintained values around 0.8 for sequences up to 700 residues, along with high designability as measured by TM-scores through joint refinement of sequence and structure(Fig. 2A,B). For binder design (Fig. 2C), Protein Hunter combined with Boltz-2 achieved higher ipTM scores in 9 of 11 targets, and with Chai in 7 of 11 targets, compared to RFdiffusion, while maintaining strong pLDDT values. Furthermore, the method enables binder design for multimeric proteins (Fig. 3A), generating binders on TNF-*α* trimer surfaces without disrupting inter-trimer interactions or mispositioning them within the protein core.

**Figure 2:**
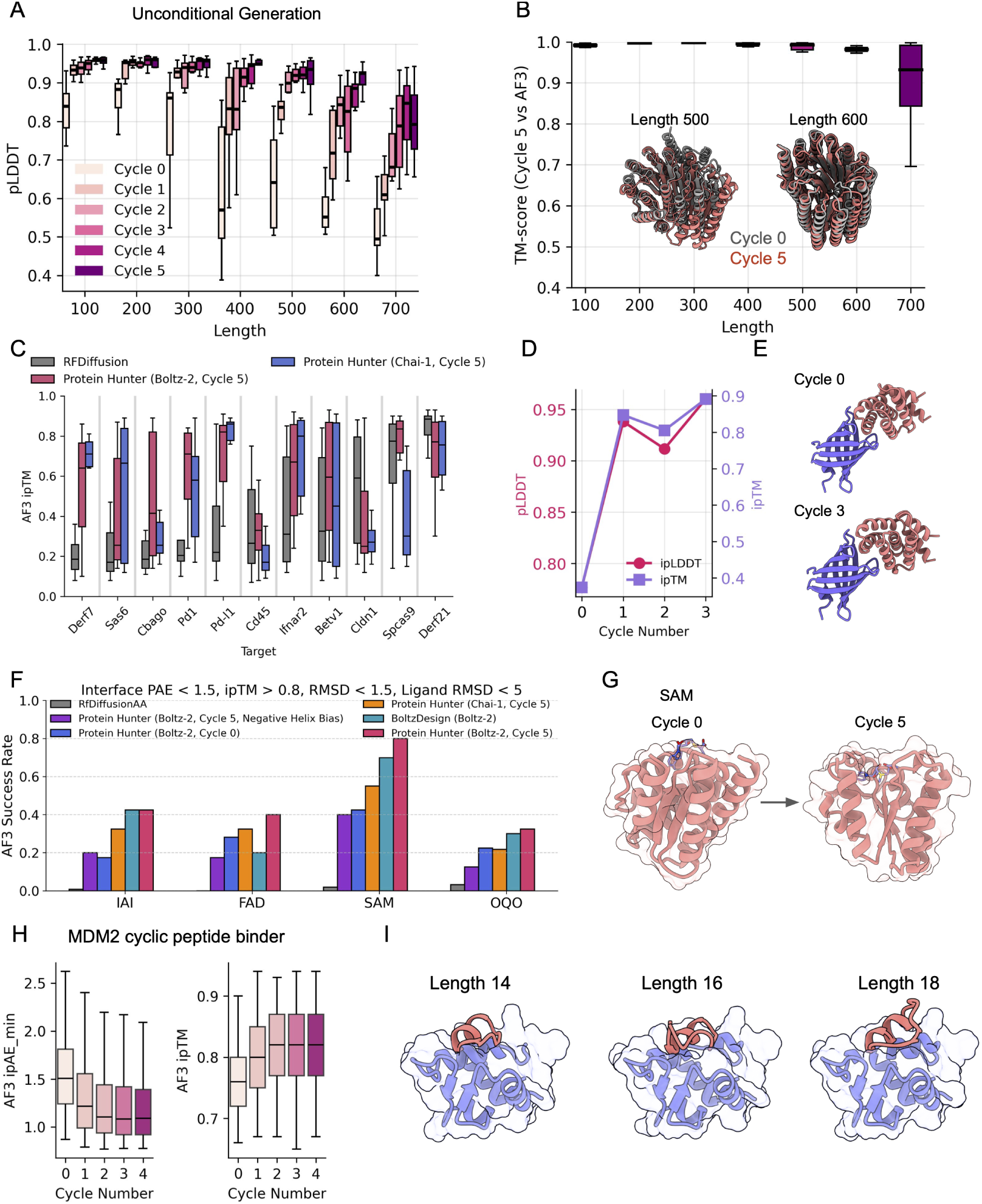
Protein Hunter cycles structure prediction and sequence design for confident protein generation. (A) Unconditional generation for 100–700 residue proteins using Protein Hunter (Boltz-2). (B) Designability measured by TM-scores between Cycle 5 structures and AF3-repredicted structures, with example unconditional generations of 500- and 600-residue proteins. (C) Binder design on protein targets: Protein Hunter vs. RFdiffusion. (D) BBF-14 binder design showing pLDDT and ipTM across cycles. (E) BBF-14 binder structures at cycle 0 and cycle 3. (F) Small-molecule binder success rates measured by AF3 (40 designs per ligand via MPNN). (G) SAM small-molecule binder designs from cycle 0 to cycle 5. (H) MDM2 protein-binding cyclic peptide designs, showing AF3 iPAE(min) and ipTM scores from cycles 0 to 4. (I) Examples of designed peptides of varying lengths (14, 16, and 18 residues).

**Figure 3:**
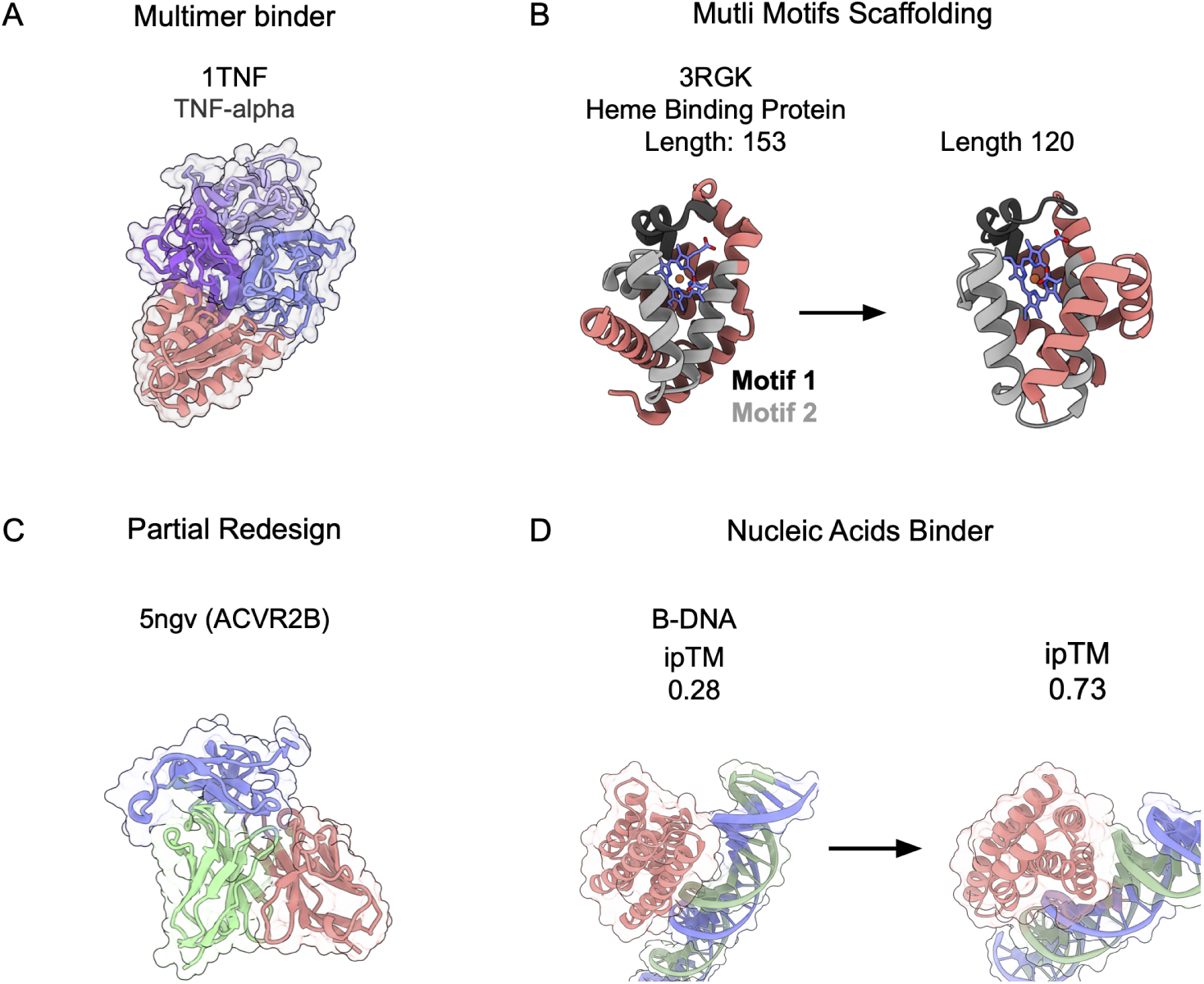
Protein Hunter designs binders for diverse biomolecules and supports motif scaffolding and partial redesign. (A)TNF-*α* trimer binder. (B) Heme-binding protein: motifs 1 and 2 provided, remaining scaffold redesigned (left: 3RGK; right: length 120). (C) Partial redesign of 5NGV antibody light- and heavy-chain CDRs. (D) Nucleic acids targeting a B-DNA, where iterative cycling improved iPTM from 0.28 to 0.73.

Protein Hunter also enables all-atom binder design. Using the same iterative approach on four chemically diverse targets previously tested with RFdiffusion-AA and BoltzDesign1, small-molecule binders were generated and evaluated with AlphaFold 3. Success rate is defined by backbone RMSD ≤1.5 Å, ligand RMSD ≤5 Å, interface PAE ≤1.5, and ipTM ≥ 0.8. By these criteria, Protein Hunter reached higher success rates across all targets compared with RFdiffusion-AA, BoltzDesign2, and its single-pass mode (Figure 2F,G). Importantly, these results highlight that iterative cycling reduces the need for extensive sequence sampling: structure and sequence are progressively co-optimized across cycles, yielding strong backbone–sequence compatibility without requiring large-scale sampling and refolding for validation.

Lastly, Protein Hunter supports the design of protein-binding macrocyclic peptides. Cyclization is implemented through cyclic positional encoding, following established methods [Rettie et al., 2025a, Zhang et al., 2025, Rettie et al., 2025b]. Similar to unconditional and binder-specific generation, cyclic-sequence conditioning further enhances in silico success rates, as demonstrated in Figure 2H, which shows the MDM2 cyclic peptide binder design results, and in the varied-length peptide examples shown in Figure 2I.

### 4.3 Protein Hunter Generalizes to Diverse Protein Design Tasks

Protein Hunter extends to multi-motif scaffolding, partial redesign, and binder generation for diverse molecular targets. For motif scaffolding (Fig. 3B, 13), we demonstrate a heme-binding protein in which two motifs adjacent to the heme ligand (all-atom distance *<* 5 *Å*) were fixed while diffusion hallucination rebuilt the surrounding scaffold and termini. For partial redesign (Fig. 3C, 14), we applied the method to the 5NGV antibody structure, retaining framework regions of the light and heavy chains while optimizing the CDRs [Wang et al., 2025]. Beyond proteins, the framework also designs small-molecule, DNA, and RNA binders (Fig. 3D), where successive structure–sequence cycles increased pLDDT and ipTM, improving structural accuracy and predicted interaction quality.

## 5 Discussion

Protein Hunter unites zero-shot structure prediction with iterative sequence–structure refinement, combining the speed of generative diffusion with the precision of co-optimization. Cycling reduces biases such as alanine enrichment, improves foldability, and adding pairwise bias terms to pair representations allows control of secondary structure, for example, a negative helix bias promotes *β*-sheet–rich designs and further improves the diversity of structures. The diffusion hallucination process can be further steered by incorporating potential terms such as contact and pocket constraints, enabling specific guidance without retraining. Beyond binder generation, Protein Hunter readily supports multi-motif scaffolding, partial redesign, and all-atom targets, enabling catalytic-site embedding, antibody CDR optimization, and design of small-molecule or nucleic-acids binders. Future work will focus on more accurate catalytic side-chain modeling, broader secondary-structure sampling, and better coverage of diverse fold distributions. Incorporating sparse autoencoders to extract and activate fold-specific features from pair representations may further expand structural diversity. Aligning with developments in structure prediction models and strategies for sampling structures, Protein Hunter will be able to generate an even wider range of functional proteins.

## 6 Acknowledgements

We would like to thank Katelyn Campbell for helpful suggestions. Y.C. acknowledges financial support from the SBS scholarships and the Takeda Fellowship. S.O. acknowledges funding from NSF grant MCB2032259, and Amgen. We extend our special thanks to Rohith Krishna for providing the RfDiffusion-generated backbones.

## 7 Data and code availability

All data and source code is available at https://github.com/yehlincho/Protein-Hunter

## A Appendix

### A.1 Diffusion Hallucination from Single–Amino-Acid Sequences

To identify which single–amino-acid sequence best matches the natural protein distribution, we modeled background residue–distance statistics using the CATH database Orengo et al. [1997], Anishchenko et al. [2021]. For each residue pair (*i, j*) we defined the absolute sequence separation

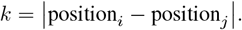

Observed C_*β*_–C_*β*_ distances *d* for each *k* were binned into a 64-bin distogram spanning 2–22 Å. The background probability

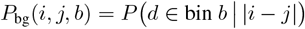

gives the likelihood that the distance between residues *i* and *j* falls into bin *b*, with

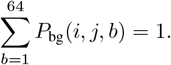

We generated 120-residue sequences composed of a single repeated amino acid (e.g., (aa)_120_) and supplied them to the model in single-sequence mode. After three recycling steps, we extracted pairwise feature representations from the Boltz Pairformer trunk and computed their Spearman correlation with the background distribution. The resulting distogram maps are shown in Fig. 4, and an example of single–amino-acid sequence hallucination targeting CLDN1 is shown in Fig. 5.

**Figure 4:**
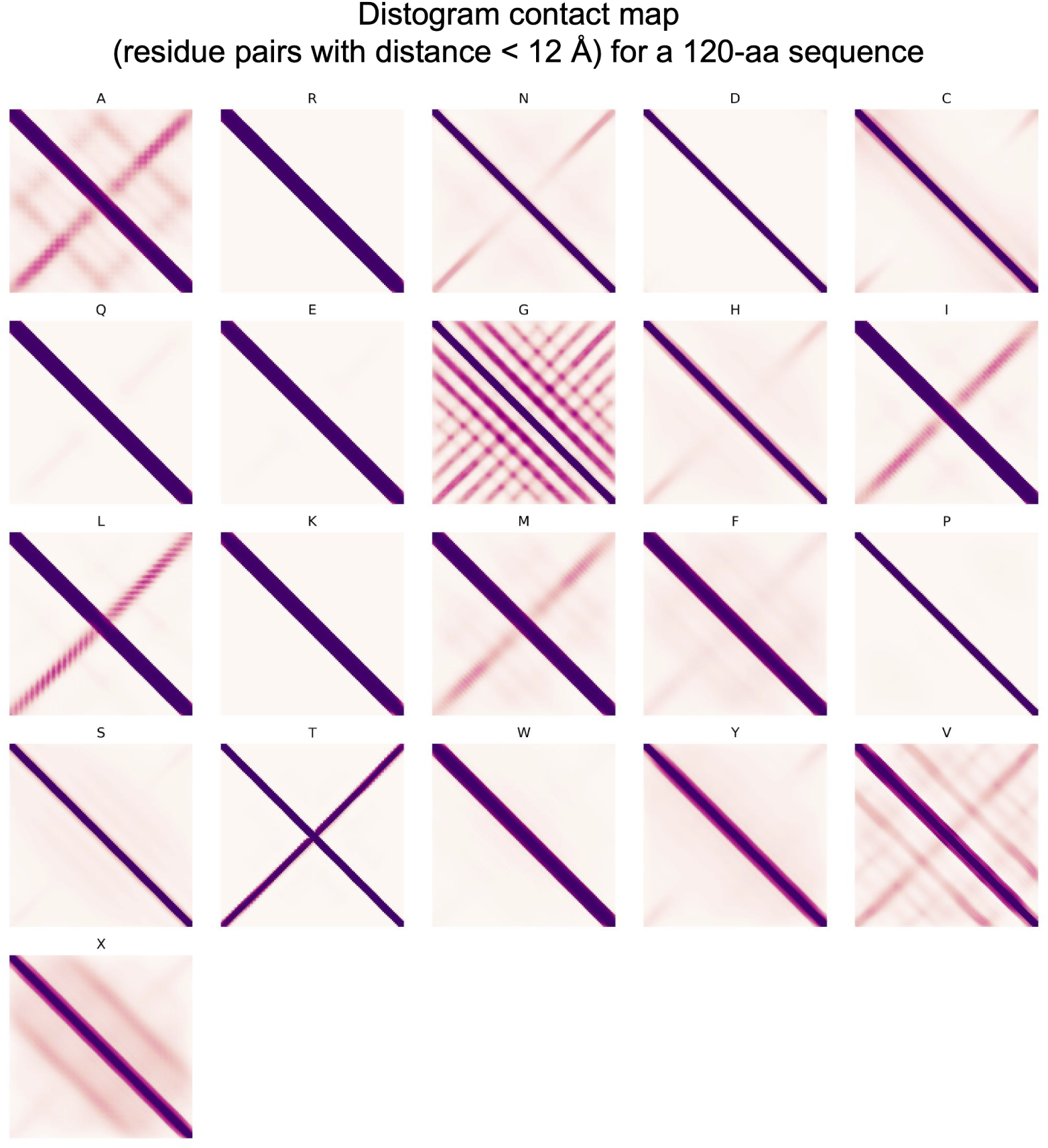
Distogram contact map of single–amino-acid sequences, obtained by summing all distance bins *<*12 Å. Shown is a 120-residue sequence tested for each of the 20 standard amino acids and for the unknown token “X.”

**Figure 5:**
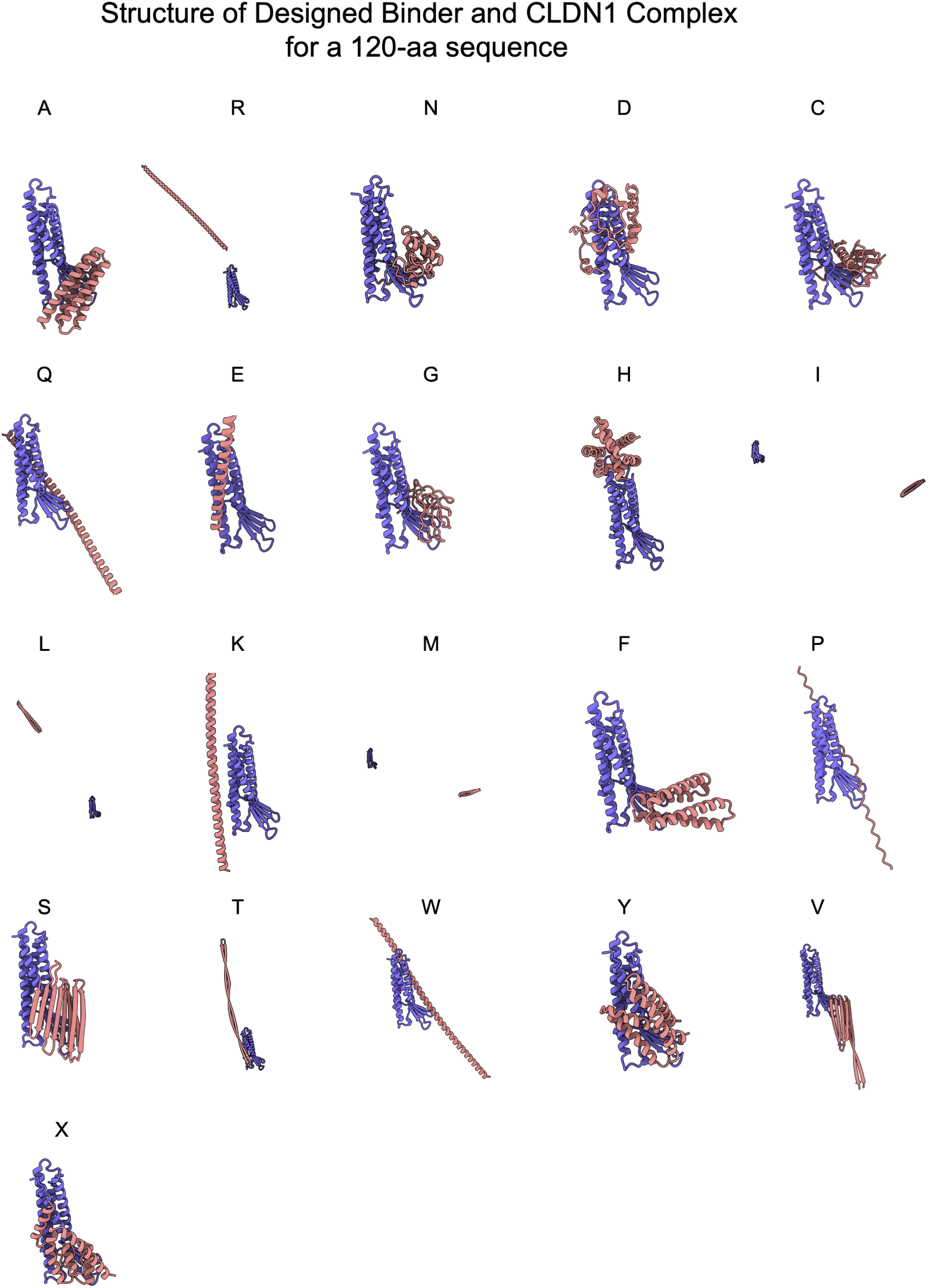
Predicted binder design using CLDN1 as the target. A 120-residue sequence was tested for each of the 20 standard amino acids. The target structure is shown in purple and the predicted binder structure is shown in pink.

**Figure 6:**
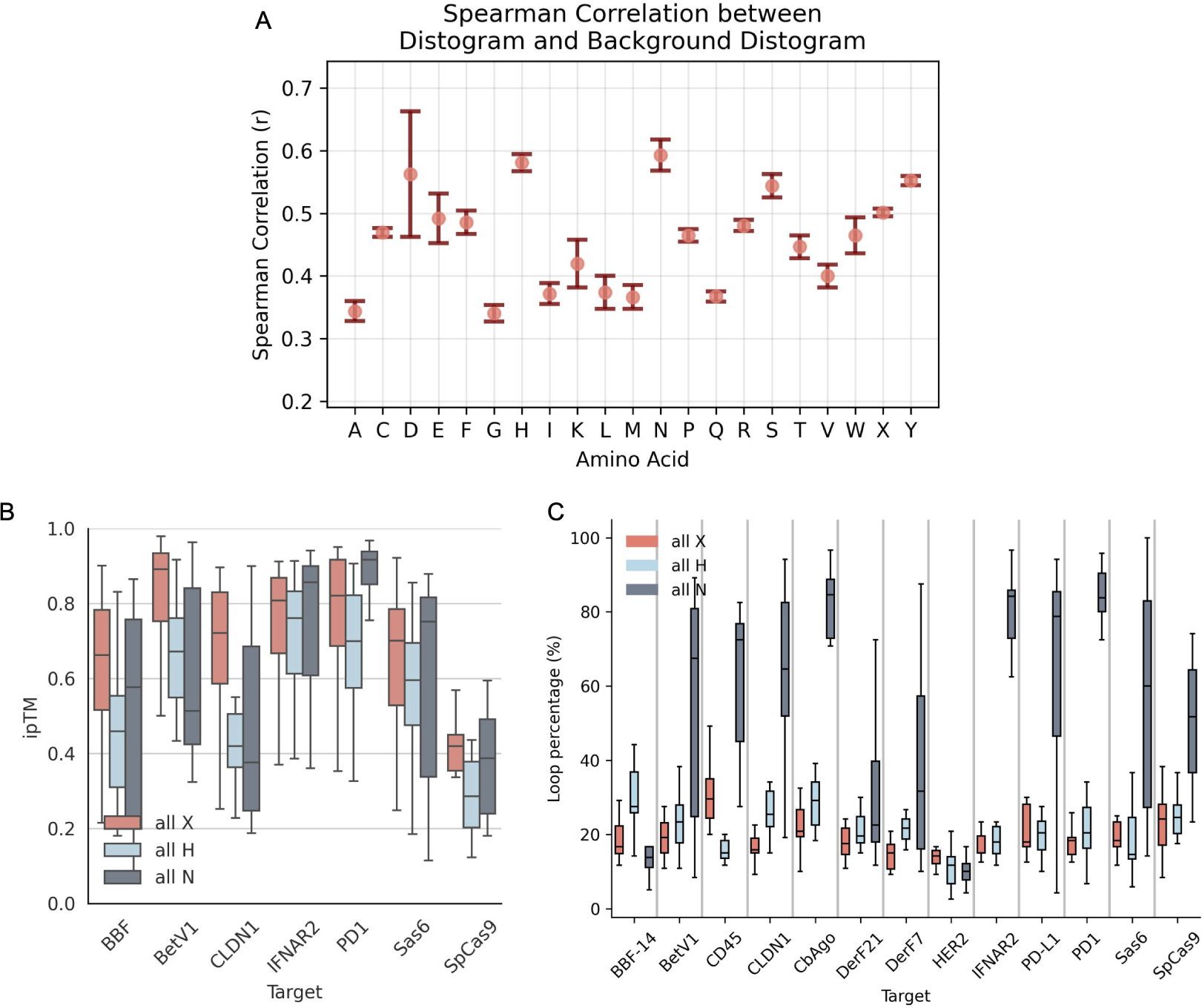
(A) Spearman correlation between single–amino-acid sequences and background distograms extracted from CATH domains. (B) Binder design starting from hallucinations using all “X,” all “H,” or all “N” sequences, followed by ProteinMPNN redesign, with performance measured by Af3 ipTM. (C) Loop content (percentage of residues in loops) of the designed binders.

**Figure 7:**
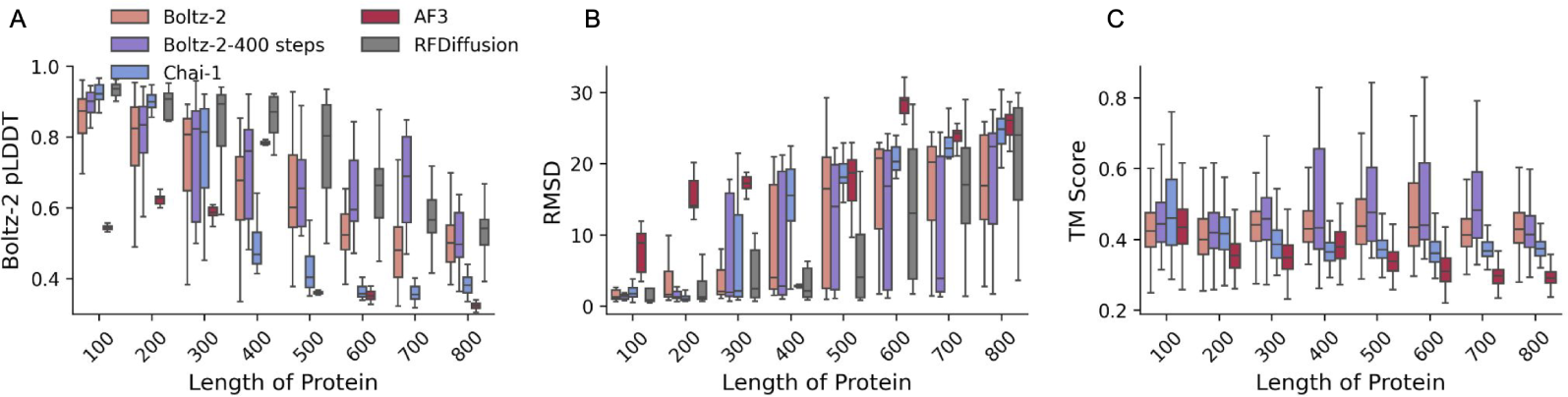
Unconditional zero-shot generation of protein structures using different AF3-style models. After zero-shot structure generation, designs were redesigned with SolubleMPNN and refolded using the Boltz-2 model in single-sequence mode. (A) pLDDT scores from Boltz-2; (B) RMSD between generated and refolded structures; (C) average pairwise TM-scores across generated structures, used to measure structural diversity, comparing Boltz-2 (200 diffusion steps), Boltz-2 (400 diffusion steps), Chai-1 (ESM embedding), AF3, and RFdiffusion as baselines.

**Figure 8:**
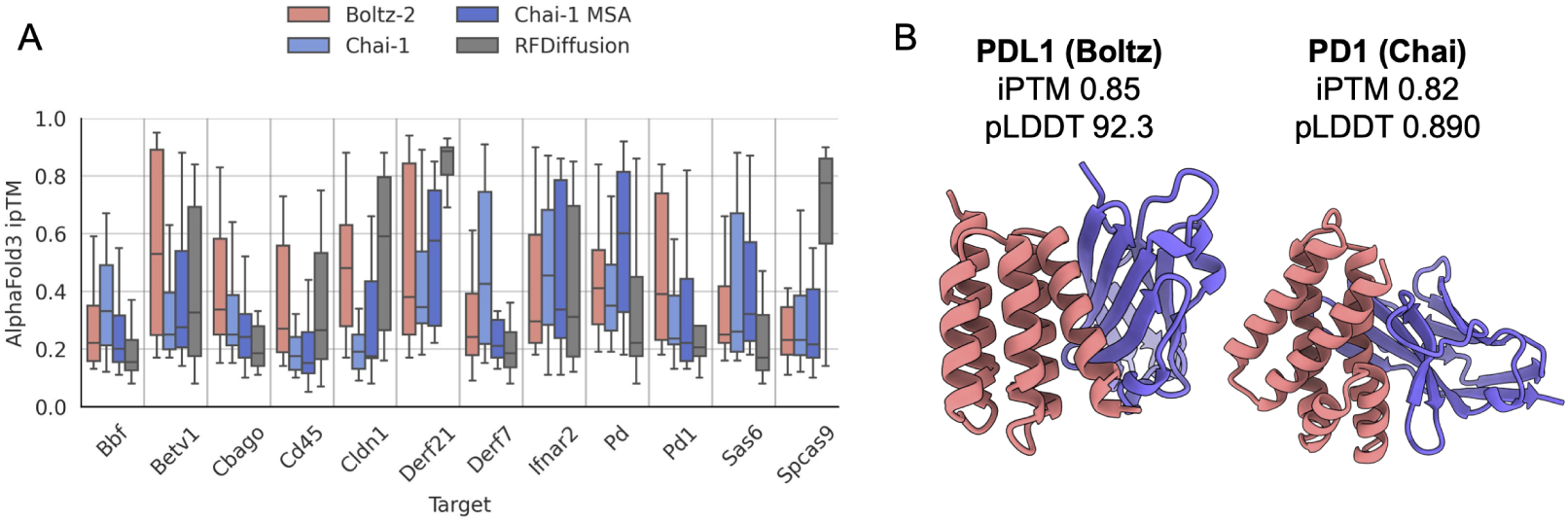
(A) Cycle 0 structures generated from an all “X” sequence, redesigned with ProteinMPNN, and evaluated by ipTM. Results are compared across Chai-1 ESM embeddings only, Chai-1 ESM embeddings with MSA, and Boltz-2, using RFdiffusion as the baseline. (B) Successful one-shot designs from Boltz and Chai models showing high ipTM and pLDDT scores against PDL1 and PD1 targets.

**Figure 9:**
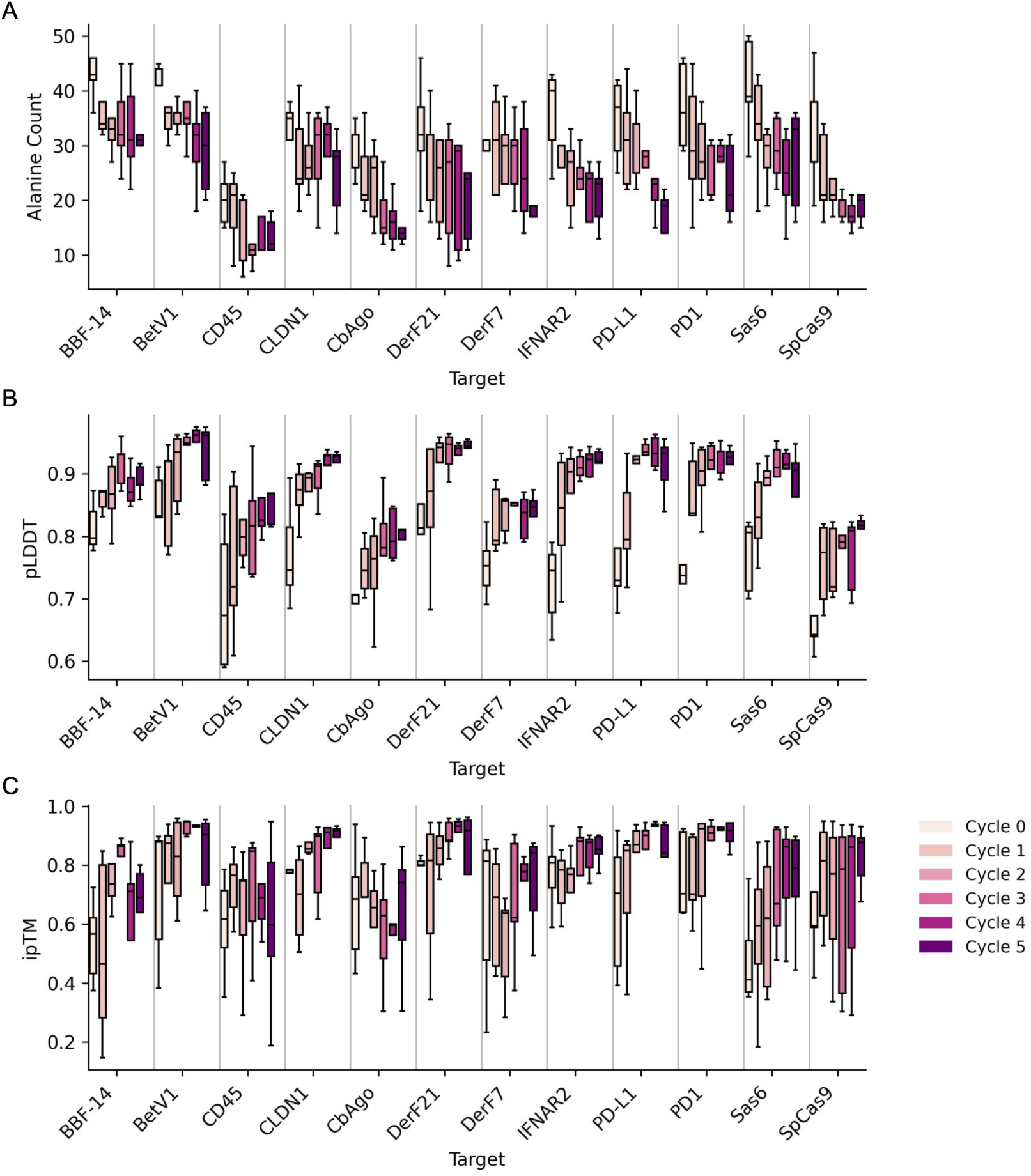
Cycling effects of the Boltz-2 model on binder generation toward protein targets from cycle 0 to cycle 5. (A) Alanine count. (B) pLDDT. (C) ipTM.

**Figure 10:**
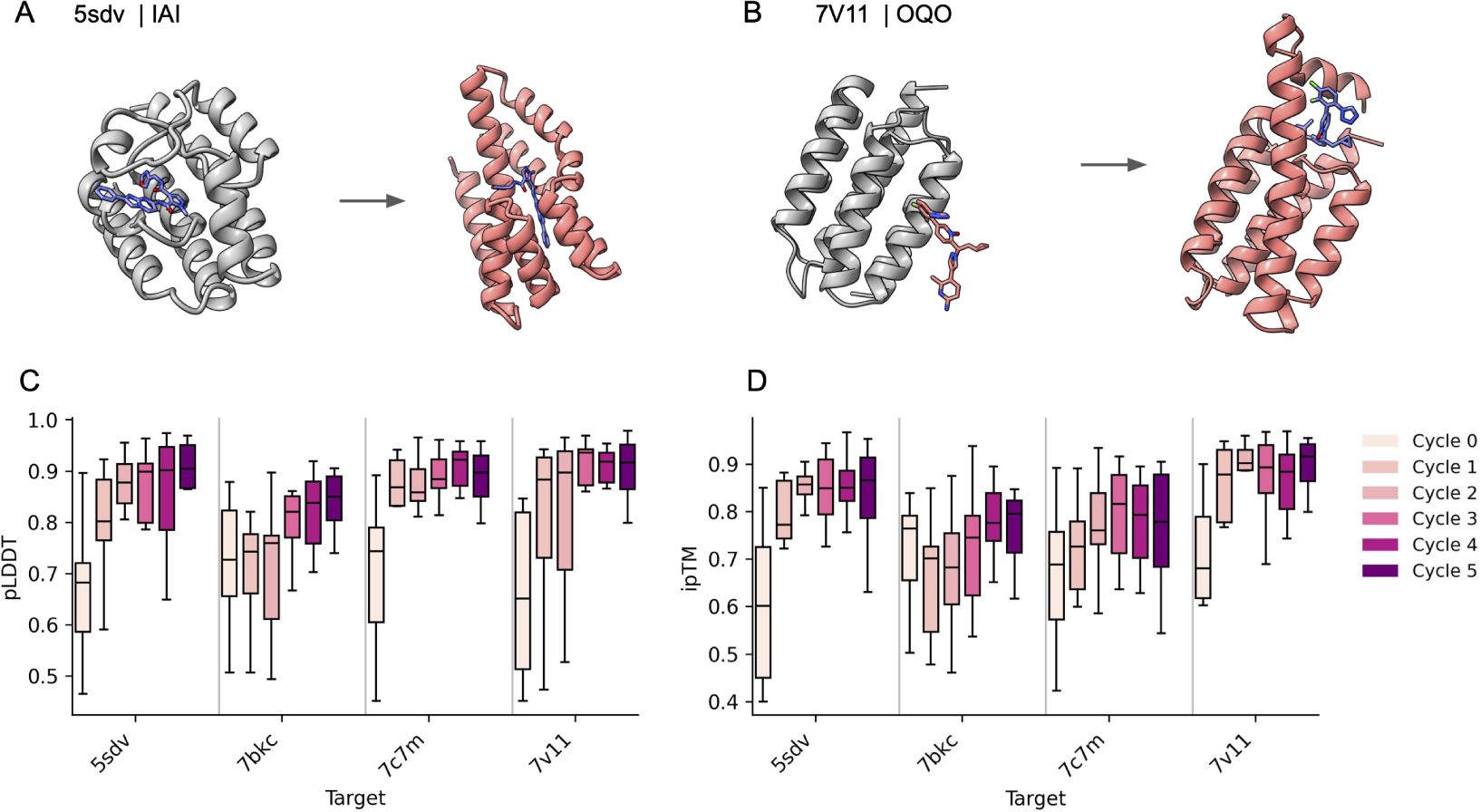
Cycling effects on small-molecule binder generation. (A) Binder design for the IAI small molecule. (B) OQO cycling improves proximity of the small molecule to the binder. (C) pLDDT and (D) ipTM scores of designed binders across four small-molecule targets during cycling.

**Figure 11:**
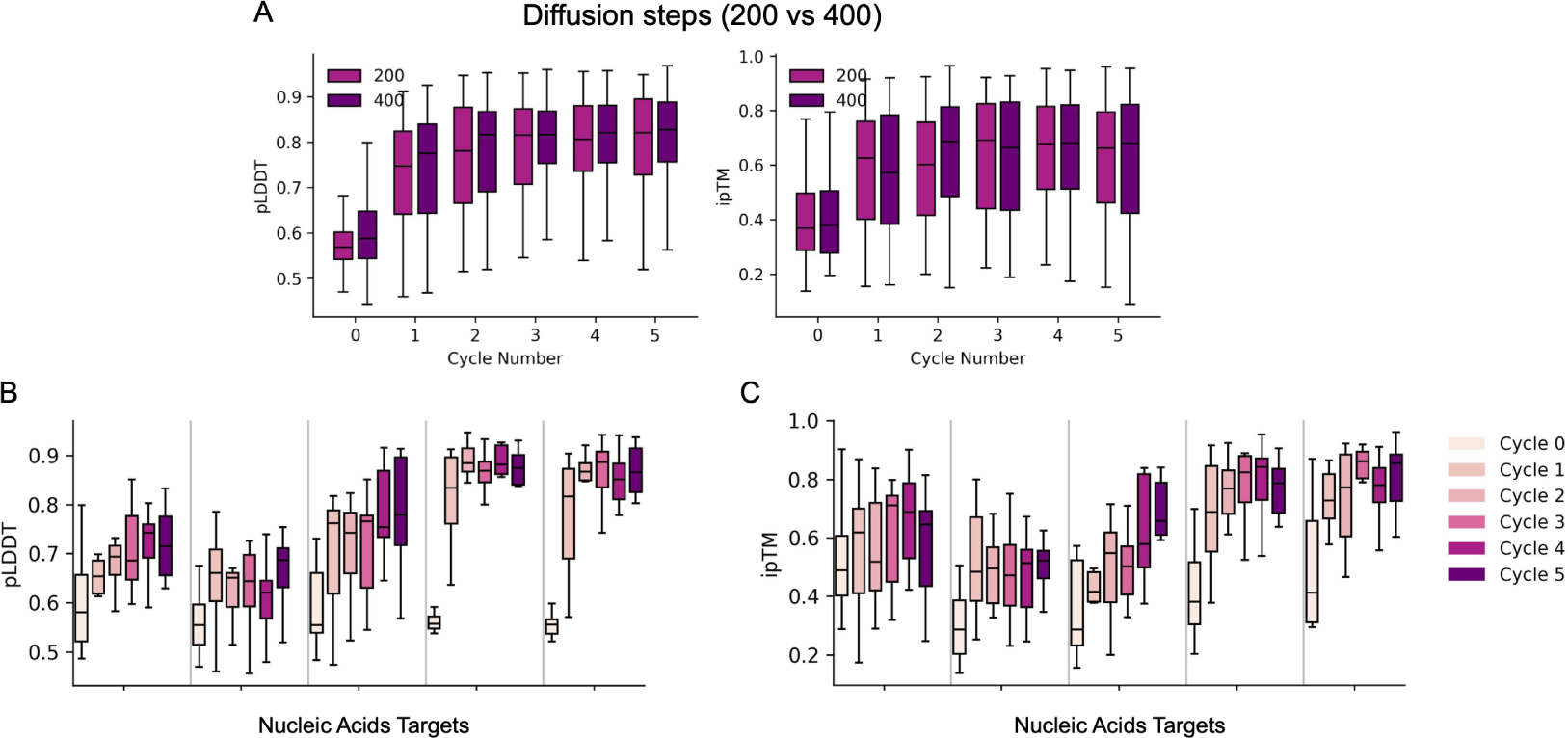
Nucleic-acids binder generation. (A) Effect of varying the number of diffusion steps on pLDDT and ipTM, showing no major differences. (B) pLDDT and (C) ipTM of nucleic-acids binders during cycling across five targets.

**Figure 12:**
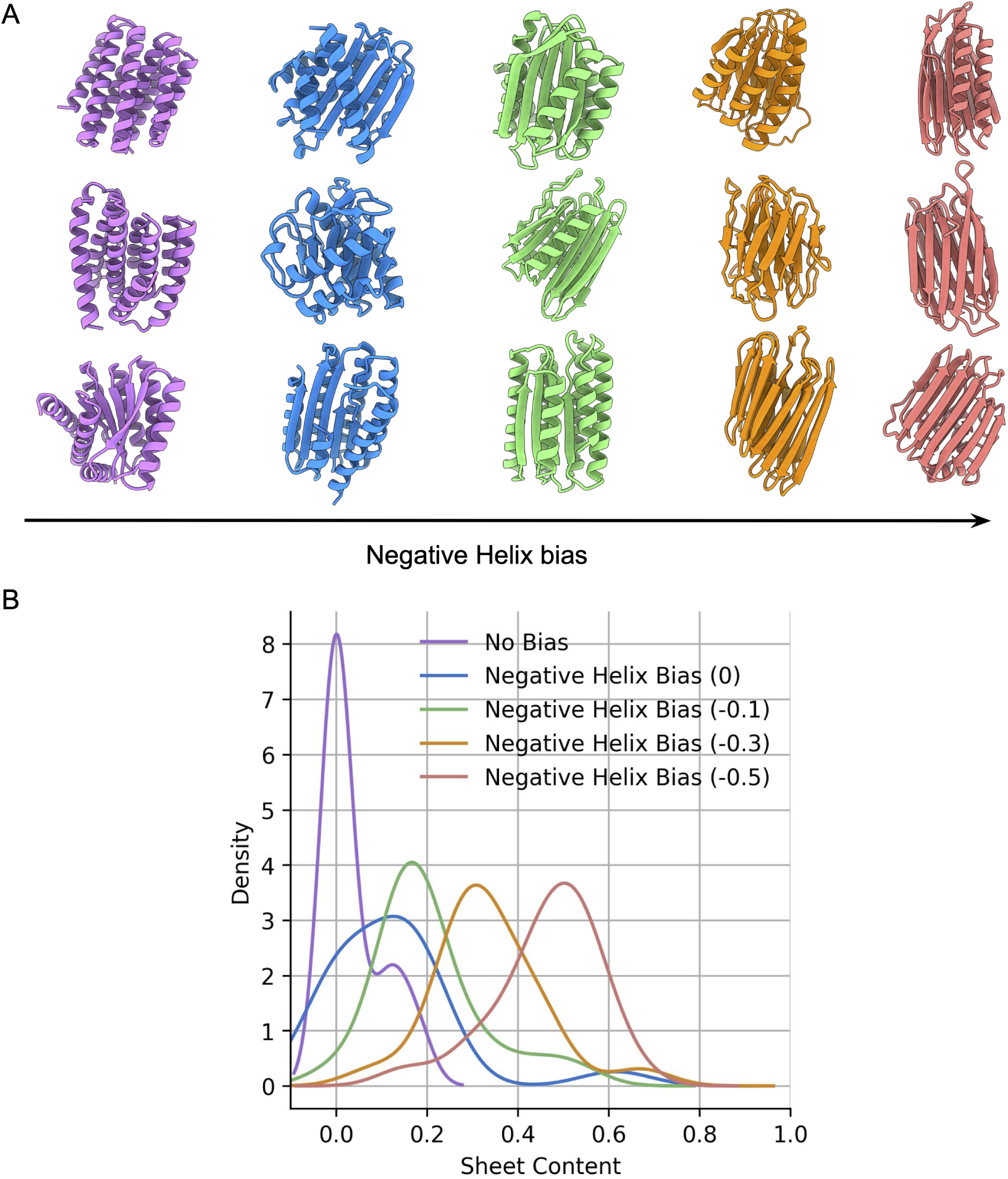
Introducing a negative helix bias in the pair features before adding them to the diffusion model decreases helix content and increases sheet content. (A) Examples of structures show increasing sheet content as the negative helix bias increases. All structures are generated with a cycle 0 setting. (B) The density plot shows the sheet fraction of designs across different bias values.

**Figure 13:**
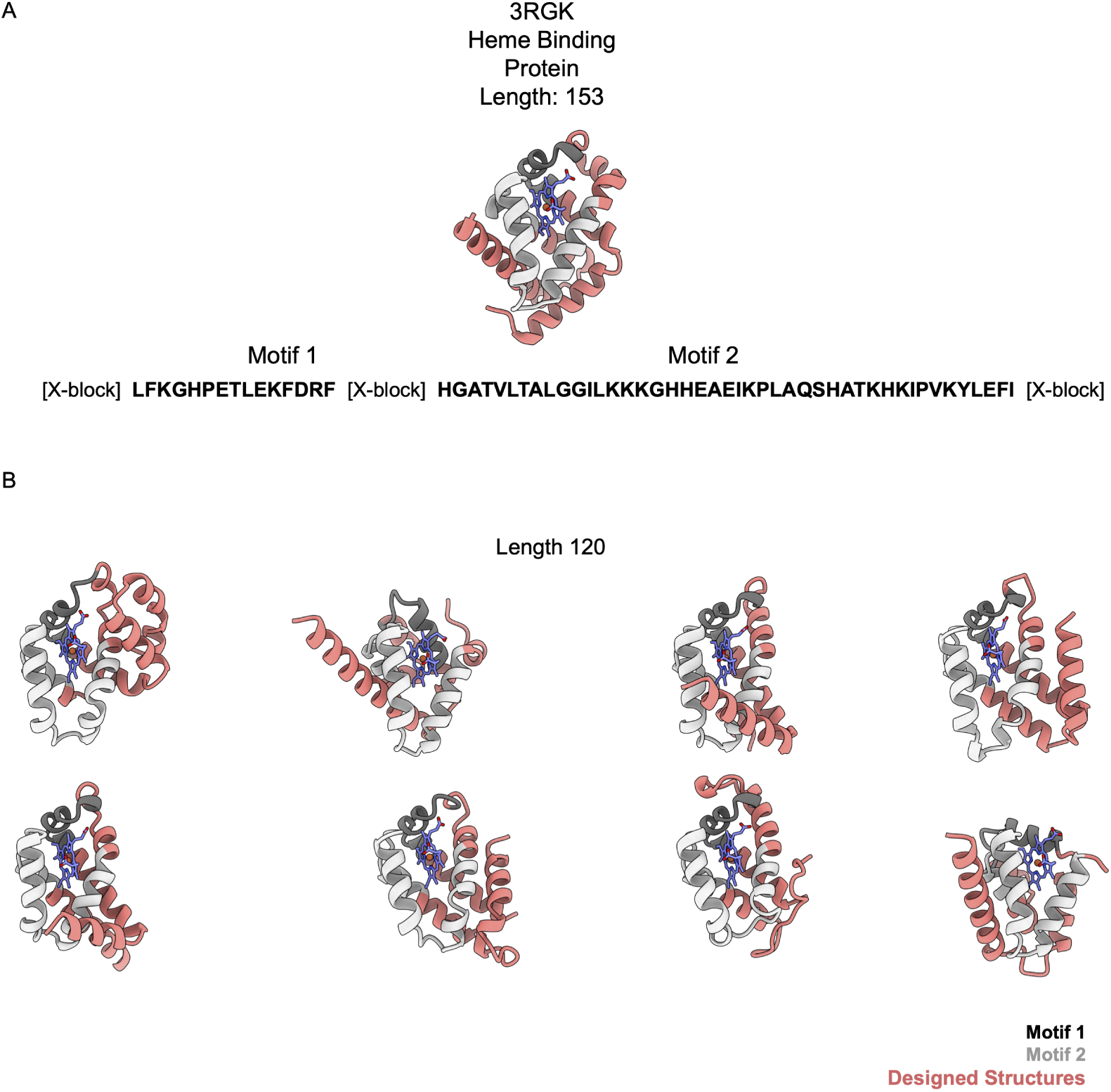
Multi-motif scaffolding. (A) Given two adjacent heme-binding motifs (1 and 2) and their co-ordinates as template features, all other positions were filled with random “X” tokens. Motif locations and inter-motif gaps were randomly chosen, and the model sampled multiple heme-binding protein designs. Gray indicates motif regions and pink indicates redesigned regions. (B) Examples of heme-binding proteins generated from multi-motif scaffolding.

**Figure 14:**
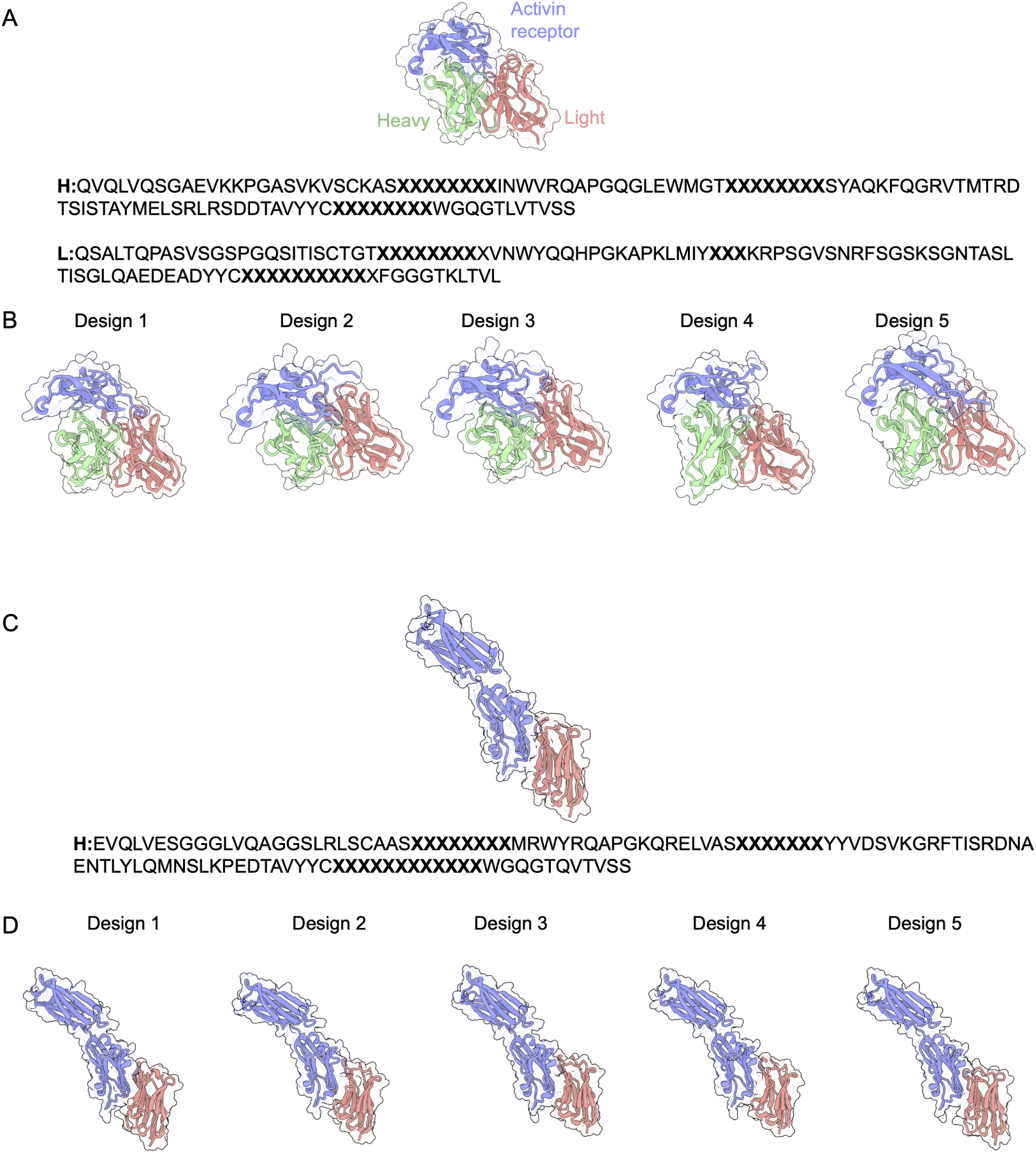
Partial Redesign. (A) Activin receptor-targeting heavy-light chain antibody (PDB: 5ngv). Purple: Activin receptor; green: heavy chain; pink: light chain. (B) Five CDR-redesigned heavy and light chain designs. (C) VHH binder complex with PD-L1 (PDB: 8aom). Purple: PD-L1; pink: VHH. (D) Five CDR-redesigned VHH designs.

**Figure 15:**
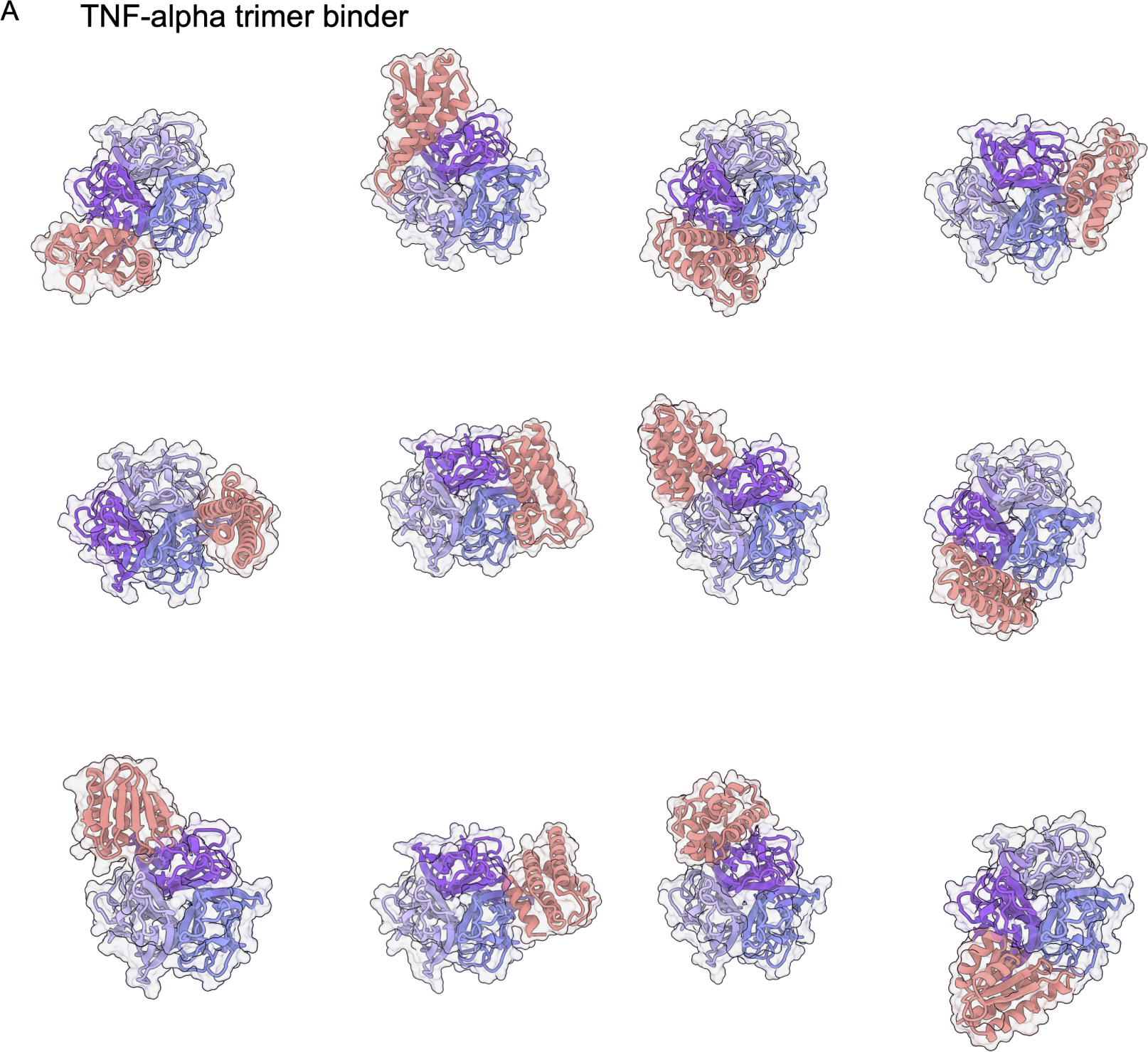
Multimeric protein binder design. (A) TNF-*α* trimer binding protein generation. Purple: trimer; pink: binder.

## Footnotes

2 RFdiffusion was also evaluated with single-sequence refolding for comparability, which may yield success rates different from those originally reported.

